# Apolipoprotein B and Interleukin 1 Receptor Antagonist: Reversing the Risk of Coronary Heart Disease

**DOI:** 10.1101/2023.04.19.537591

**Authors:** Fangkun Yang, Ning Huangfu, Jiaxi Shen, Pengpeng Su, Lujie Zhu, Hanbin Cui, Shuai Yuan

**Affiliations:** Department of Cardiology, First Affiliated Hospital of Ningbo University (Ningbo First Hospital), School of Medicine, Ningbo University, Ningbo, China; Key Laboratory of Precision Medicine for Atherosclerotic Diseases of Zhejiang Province, Ningbo, China; Cardiovascular Disease Clinical Medical Research Center of Ningbo, Zhejiang China; School of Medicine, Wenzhou Medical University, Wenzhou, China; Unit of Cardiovascular and Nutritional Epidemiology, Institute of Environmental Medicine, Karolinska Institutet, Stockholm, Sweden

**Keywords:** apolipoprotein B, interleukin 1 receptor antagonist, coronary heart disease, myocardial infarction, lifestyle modification

## Abstract

**Background:** Epidemiological evidence for the link of interleukin 1 (IL-1) and its inhibition with coronary heart disease (CHD) remains controversial. We aim to investigate the cardiovascular effects of IL-1 receptor antagonist (IL-1Ra) and underlying mechanisms, as well as the potential interaction with lifestyle factors.

**Methods:** A comprehensive multivariable Mendelian randomization study was performed. Genetic variants identified from a genome-wide association study involving 30,931 individuals were used as instrumental variables for the serum IL-1Ra concentrations. Genetic associations with CHD (60,801 cases and 123,504 controls) were extracted from the CARDIoGRAMplusC4D consortium. Inverse-variance weighted method was utilized to derive effect estimates, while supplementary analyses employing various statistical approaches.

**Results:** Genetically determined IL-1Ra level was associated with increased risk of CHD (odds ratio (OR), 1.07; 95% CI: 1.03-1.17) and myocardial infarction (OR, 1.13; 95% CI: 1.04-1.21). The main results remained consistent in supplementary analyses. Besides, IL-1Ra was associated with circulating levels of various lipoprotein lipids, apolipoproteins and fasting glucose. Interestingly, observed association pattern with CHD was reversed when adjusting for apolipoprotein B (OR, 0.84; 95%CI: 0.71-0.99) and slightly attenuated on accounting for other cardiometabolic risk factors. Appropriate lifestyle intervention was found to lower IL-1Ra concentration and mitigate the heightened CHD risk it posed.

**Conclusions:** Apolipoprotein B represents the key driver, and a potential target for reversal of the causal link between serum IL-1Ra and increased risk of CHD/MI. The combined therapy involving IL-1 inhibition and lipid-modifying treatment aimed at apolipoprotein B merit further exploration.

## Introduction

Coronary heart disease (CHD) represents the leading cause of morbidity and mortality worldwide, posing a significant public health concern.^1^ Recent research has characterized CHD as a chronic inflammatory disease with inflammation-mediated atherosclerosis being a key contributing factor, which sparks interests in the potential benefits of targeting inflammation in cardiovascular diseases (CVDs).^2^ Interleukin 1 (IL-1), a prototypical proinflammatory cytokine, plays a pivotal role in the innate immune response by activating the IL-1 receptor and thus leading to the activation of downstream inflammatory mediators. IL-1 pathway has emerged as a promising therapeutic target.^3^ Nevertheless, the question remains whether targeting IL-1 related inflammation could translate into clinical benefit against cardiovascular events.

IL-1 receptor antagonist (IL-1Ra) provides natural inhibition by competing with IL-1α and IL-1β for binding to the receptor.^4^ Despite its inhibitory role, the epidemiological evidence remains controversial. A meta-analysis comprising six population-based cohorts has revealed that levels of serum IL-1Ra exhibit a positive correlation with the risk of developing CVDs.^5^ Another cross-sectional study involving 430 patients suffering from rheumatoid arthritis (RA) demonstrated a significant association between serum IL1-Ra levels and various cardiovascular comorbidities, including dyslipidemia, beta cell function, and obesity.^6^ The causality of this association remains shrouded in uncertainty, primarily due to the influence of confounding factors and the potential for reverse causation.

Several drugs targeting IL-1 have been investigated for the treatment of CVDs, including Canakinumab, Rilonacept, and Anakinra (a human recombinant IL1-Ra and a first-line agent for RA).^7^ In a randomized controlled trial (RCT) involving 80 patients with RA, Anakinra was found to significantly improve various vascular function indices as well as myocardial deformation and twisting.^8^ However, another RCT of 182 patients with non-ST elevation acute coronary syndrome suggested that treatment with Anakinra increased the risk of major adverse cardiovascular events at one year of follow-up.^9^ Notwithstanding, the duration and statistical power of previous trails have fallen short in evaluating the effect on cardiovascular outcomes.

An alternative strategy focusing on the genetic variants responsible for the inhibition of IL-1 could strengthen the causal inference.^10^ Utilizing genetic variants randomly assigned at conception as an instrumental variable (IV), it is possible to derive estimates that are less susceptible to environmental confounding factors, measurement error, and reverse causality.^11^ Our previous study has indicated the causal effect of lifetime exposure to elevated levels of IL-1Ra on an increased CHD risk, but the underlying mechanisms were not fully elucidated.^12^ Another early Mendelian randomization (MR) study suggested that long-term dual IL-1α/β inhibition increased concentrations of LDL-cholesterol, but not apolipoprotein or glycaemic traits.^13^ In recent years, the genetic determinants of IL-1Ra and cardiometabolic risk factors have been further well-characterized, which provides a promising tool to assess the causal network. In addition to pharmacological interventions, lifestyle modification represents a crucial aspect of clinical management. An investigation into the interplay between various lifestyle factors and the levels of IL-1Ra could provide insights into the heightened risk of developing CHD.

Thus, we utilized a genetic approach mimic the causal effects of serum IL-1Ra on CHD risk as well as circulating lipoprotein lipids, apolipoproteins, glycaemic traits, and blood pressure to gain insights into underlying mechanisms. Furthermore, the causal associations of obesity and lifestyle factors with IL-1Ra concentrations were examined.

## Methods

### Study Design

A comprehensive MR study was designed to investigate the causal role of IL-1Ra (Figure 1). First, we selected genetic variants as an IV for serum concentrations of IL-1Ra. Second, the combined effects of such genetic variants on RA and circulating concentrations of C-reactive protein (CRP) were assessed to examine whether the genetic IVs were valid and robustly linked with IL-1Ra concentrations. Then, the causal associations of genetically predicted IL-1Ra concentrations with the risk of CHD and MI were investigated. Third, the exploratory analysis of the genetic IVs in relation to main cardiometabolic risk factors were performed to better understand potential mechanisms between long-term pharmacological IL-1 inhibition and CHD/MI risk. Fourth, based on the above analysis, the supplementary analyses were further conducted with the adjustment of risk factor that was closely linked with IL-1Ra concentrations. Fifth, we assessed the effects of common lifestyle factors on the serum concentrations of IL-1Ra. and compared the causal effect sizes of lifestyle factors on the risk of CHD with or without the adjustment of IL-1Ra concentrations.

**Figure 1.**
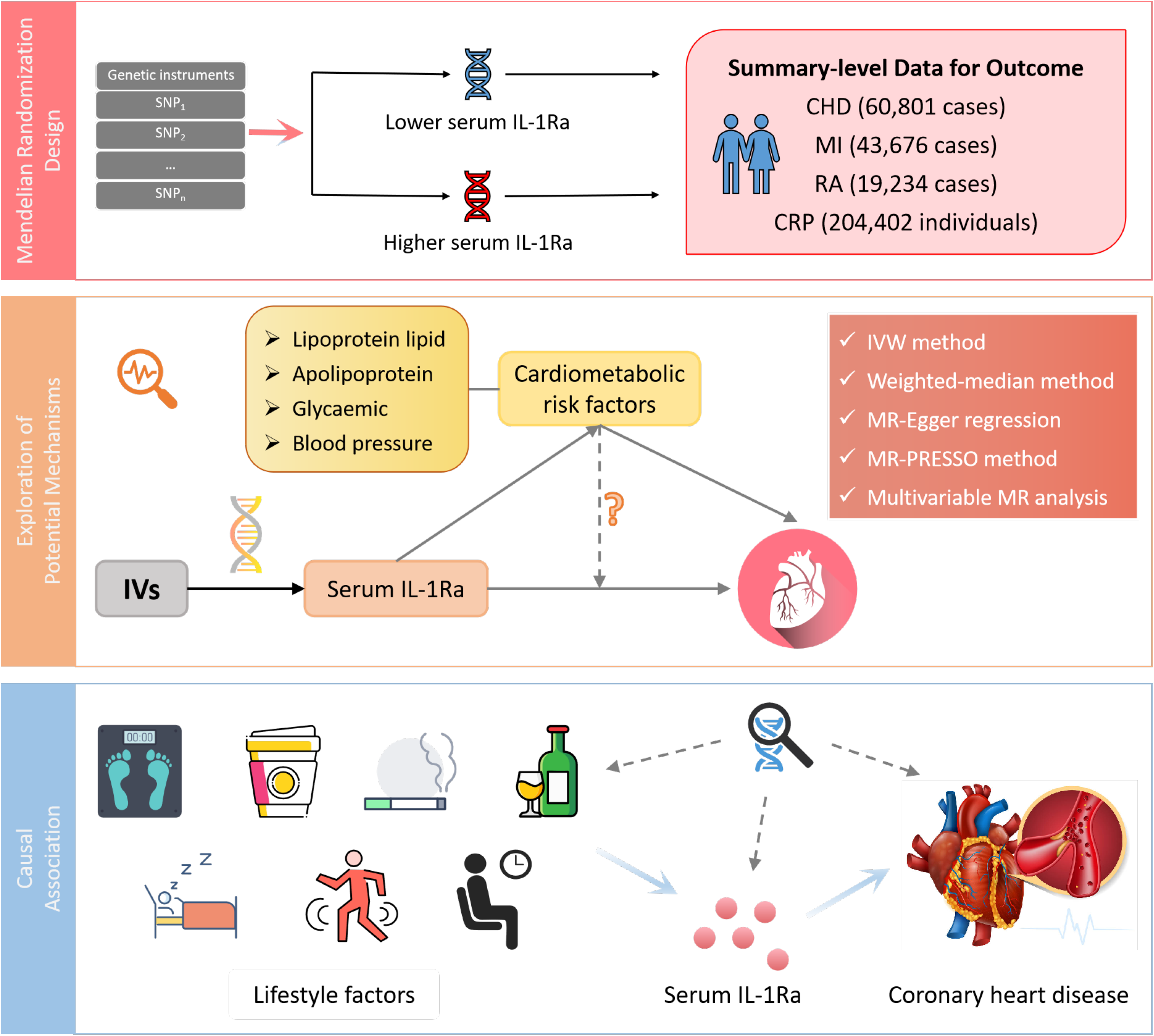
Design of the current two-sample Mendelian randomization study. IV, instrumental variable; IL1-Ra, interleukin 1 receptor antagonist; CHD, coronary heart disease; MI, myocardial infarction; RA, rheumatoid arthritis; CRP, C-reactive protein.

### Genetic Instrument Selection

Genetic variants associated with serum concentrations of IL-1Ra were identified from a genome-wide association study (GWAS) involving 30,931 individuals of European descent across 15 studies (Table 1).^14^ The primary study utilized the Olink proximity extension assay cardiovascular I panel to measure 90 cardiovascular-related proteins.^14^ Single nucleotide polymorphisms (SNPs) for serum concentrations of IL-1Ra were identified from genetic variants in the *IL1RN* gene ± 5mb region at the genome-wide significance threshold (*p* < 5×10^−8^), followed by linkage disequilibrium pruning based on the European 1000 Genomes Project reference panel (*r*^2^ < 0. 1).^15^ Among SNPs in linkage disequilibrium, the one with the smallest *p*-value was retained. Thirteen independent SNPs were selected as a genetic instrumental variable with an effect size scaled to one standard-deviations (SD) increase in IL-1Ra concentrations (Supplemental Table 1).

**Table 1.**
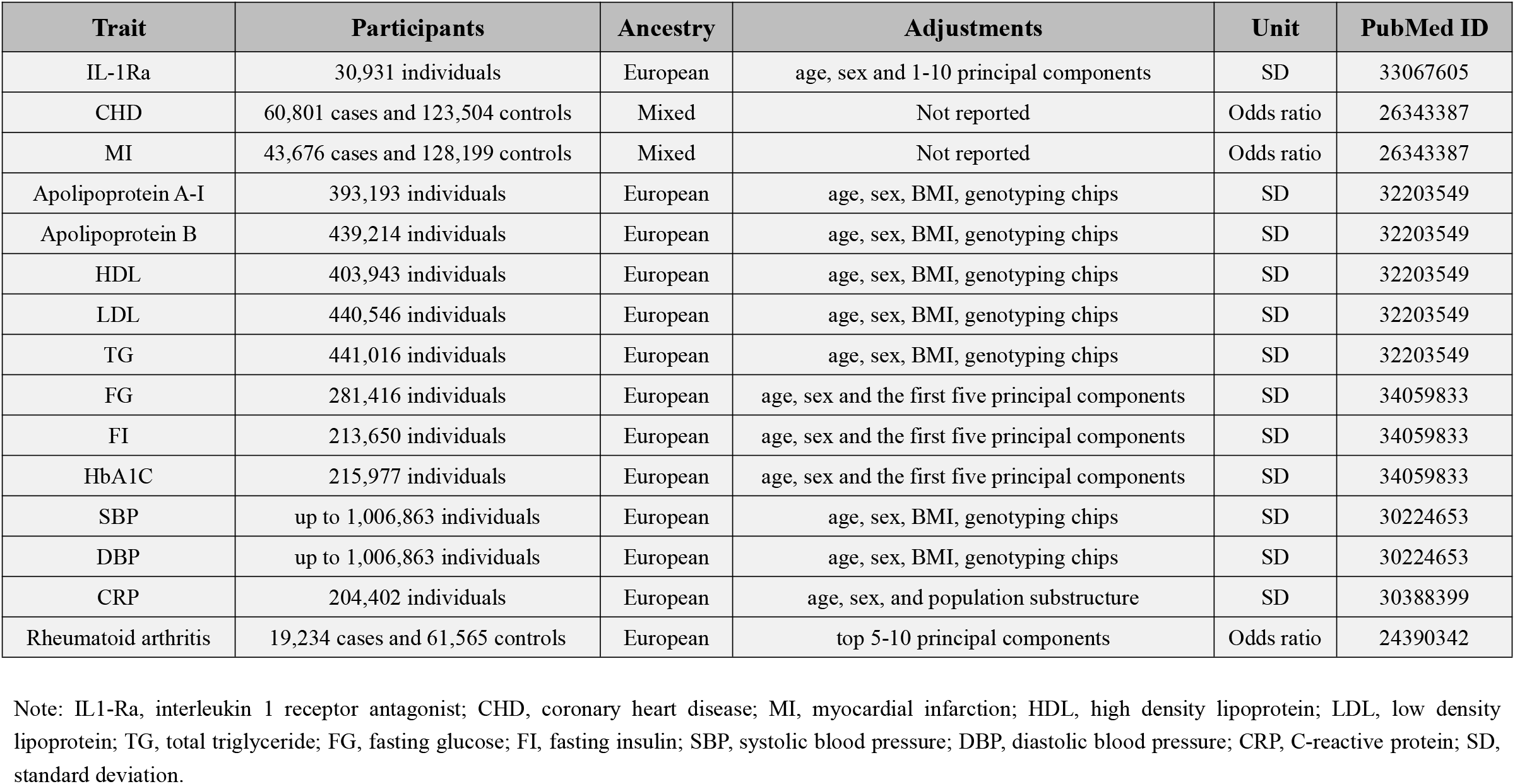
Detailed Information on Data Sources.

### Data Sources

Summary estimates of genetic associations with CHD and MI were extracted from the CARDIoGRAMplusC4D (Coronary ARtery DIsease Genome wide Replication and Meta-analysis (CARDIoGRAM) plus The Coronary Artery Disease (C4D) Genetics) consortium (Table 1).^16^ Specifically, the study included ∼185,000 participants, comprising 60,801 cases and 123,504 controls, which were from 48 different cohort studies. The CHD cases were defined as acute coronary syndrome, chronic stable angina, coronary stenosis (over 50%) or MI.^16^ The majority (∼80%) were of European ancestry and more than 70% of the total CHD cases had a history of MI.^16^ Similarly, the genetic associations with RA and CRP were obtained from corresponding GWAS.^17,18^ Detailed information were provided in the Table 1.

To explore the potential mechanisms, we obtained genetic associations with 10 main cardiometabolic risk factors, including apolipoprotein A-I, apolipoprotein B, high-density lipoprotein cholesterol (HDL-C), low-density lipoprotein cholesterol (LDL-C), and triglyceride (TG); fasting glucose (FG), fasting insulin (FI) and haemoglobin A1c (HbA1c); as well as systolic blood pressure (SBP) and diastolic blood pressure (DBP) (Table 1).^19–21^

Given the significance of lifestyle intervention in clinical management, genetic associations with various lifestyle factors, including obesity (measured by body mass index and waist circumference), smoking initiation, lifetime smoking index, alcohol drinking, alcohol dependence, coffee consumption, caffeine consumption, physical activity (both moderate-to-vigorous and vigorous), sedentary behaviour, sleep duration and insomnia were obtained from corresponding international consortia (Supplementary Table 2).^22–32^

All studies included in the GWAS had obtained ethical approval from the appropriate committees and written informed consent had been obtained from all participants. The data used in the current study were publicly available.

### Statistical Analysis

The multiplicative random-effects inverse-variance-weighted (IVW) method, as the primary statistical model, was employed to estimate and associations of IL-1Ra concentrations with the risk of CHD, MI and RA, as well as the circulating concentrations of CRP.^33^ In addition, some supplementary analyses were conducted to investigate potential horizontal pleiotropy and assess the consistency of primary results, including the weighted-median method, MR-Egger regression, and MR Pleiotropy Residual Sum and Outlier (MR-PRESSO) methods.^34–36^ The weighted median method represents a dependable approach for obtaining consistent causal estimates when up to 50% of the weight arises from the invalid instruments.^34^ The MR-Egger regression can detect the potential pleiotropy leveraging the embedded intercept test while also provide a corrected estimate accounting for the detected pleiotropy if any. However, it may entail sacrificing statistical power to some extent.^35^ On the other hand, the MR-PRESSO method can detect potential outliers and generate relatively unbiased causal estimates after removal of outliers.^36^ Furthermore, the embedded distortion test can be employed to evaluate the disparities between the obtained estimates before and after outlier removal.^36^ We also undertook the leave-one-out analysis and utilized the scatter plot to depict the associations of genetically predicted IL-1Ra concentrations with CHD/MI to examine whether a single SNP drove the association. Multivariable MR (MVMR) analysis was performed to explore the potential mechanisms and to assess whether the associations of IL-1Ra concentrations with CHD/MI changed after the adjustment for cardiometabolic risk factors.^37^ The two-step network MR mediation analysis was further conducted to investigate the extent to which any effect of IL-1Ra concentrations on CHD/MI might be mediated through apolipoprotein B.^38^ And we estimated the proportion mediated by multiplying the effect size of IL-1Ra concentrations on apolipoprotein B by that of apolipoprotein B on CHD/MI. Besides, we further investigated the causal effect of CHD and MI on IL-1Ra concentrations, considering the possibility of reverse causality. When a SNP was unavailable, we employed the SNiPa (http://snipa.helmholtz-muenchen.de/snipa3/) to search for a proxy SNP in linkage disequilibrium (*r*^2^>0.8) with the unavailable SNP based on the European population genotype data. The statistical significance threshold for the association of IL-1Ra concentrations with cardiometabolic risk factors was set at a two-sided *p* value of <0.005 (=0.05/10 tests). The statistical analyses were conducted with R software (version 4.2.0) using ‘TwoSampleMR’, ‘MendelianRandomization’, and ‘MR-PRESSO’ packages.^36,39,40^

## Results

The IVW analyses yielded compelling results demonstrating significant associations of genetic predicted higher concentrations of IL-1Ra with an increased risk of RA as well as decreased levels of CRP (Table 2). However, the corresponding risk increased by 9% and 13% per 1-SD increase in serum IL-1Ra for CHD (95%CI: 1.03-1.17; *p*=6.9×10^−3^) and MI (95%CI: 1.04-1.21; *p*=2.2×10^−3^), respectively. The sensitivity analyses indicated that MR associations of IL-1Ra with CHD/MI risk kept consistent in the majority of statistical methods, supporting the robustness of our findings, albeit with wider 95% CIs in the MR-Egger regression (Table 2; Supplementary Table 3). Neither MR-PRESSO nor MR-Egger intercept test detected potential directional pleiotropy. The results of the leave-one-out analysis suggested that the causal association between IL-1Ra concentration and increased risk of CHD/MI was not drastically driven by any individual variant (Supplementary Figure 1-6).

**Table 2.**
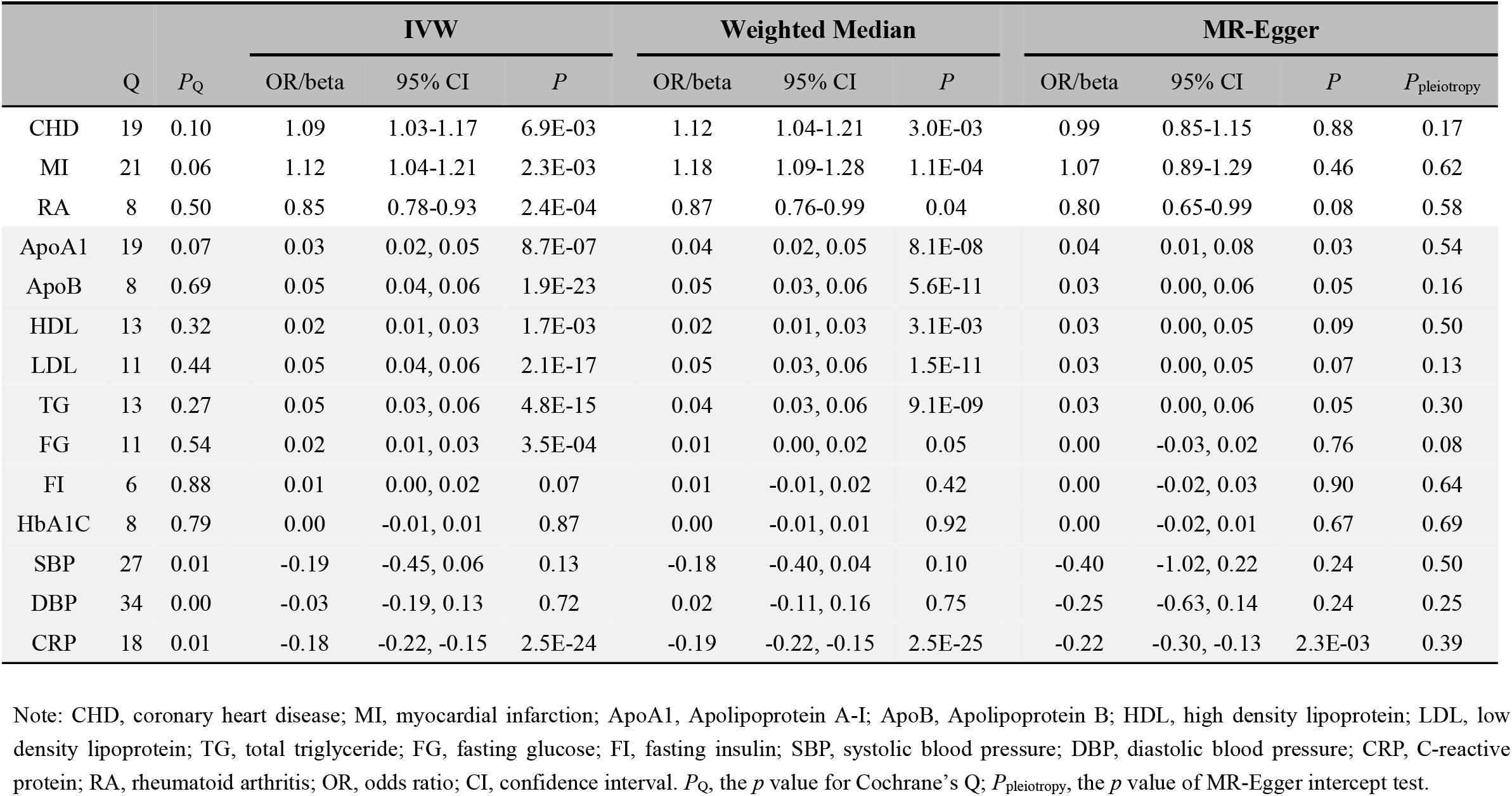
Associations of Genetically Determined IL-1Ra Level with CHD, MI and Cardiometabolic Risk Factors.

To better understand the potential mechanisms underlying the observed causal associations, we conducted analyses examining the relationship between IL-1Ra and 10 different cardiometabolic risk factors (Table 2). The genetic predisposition towards a 1 SD increase in IL-1Ra concentration was found to be significantly associated with a roughly 0.05 SD increase in apolipoprotein B, LDL and TG concentration. Additionally, apolipoprotein A-I, HDL and FG were also closely linked with IL-1Ra concentration. However, no evidence was found to support the association of IL-1Ra concentration with blood pressure. We then reassessed the association between IL-1Ra and CAD/MI with adjustment for genetically predicted cardiometabolic traits (Figure 2). In the MVMR with adjustment for apolipoprotein A-I, LDL, HDL and FG, the effect estimates were slightly attenuated, compared to the primary results without adjustments. Interestingly, we found that the association pattern between IL-1Ra concentration and the risk of CHD/MI drastically changed on accounting for apolipoprotein B concentrations. The direction of causal effect was reversed. After controlling for apolipoprotein B levels, genetically predicted higher IL-1Ra concentrations were associated with a reduced risk of CHD (OR, 0.84; 95%CI: 0.71-0.99) and MI (OR, 0.85; 95%CI: 0.71-1.02). The two-step mediation analysis indicated that 29% (95%CI: 21% to 37%) and 20% (95%CI: 14% to 27%) of the detrimental effect of serum IL-1Ra on CHD and MI was mediated through apolipoprotein B concentrations, respectively (Figure 3). Reverse-direction analyses were also performed to assess the potential reverse causality, and no consistent evidence was found for the reverse causal effect of CHD or MI on the serum IL-1Ra level (Supplementary Table 4-7).

**Figure 2.**
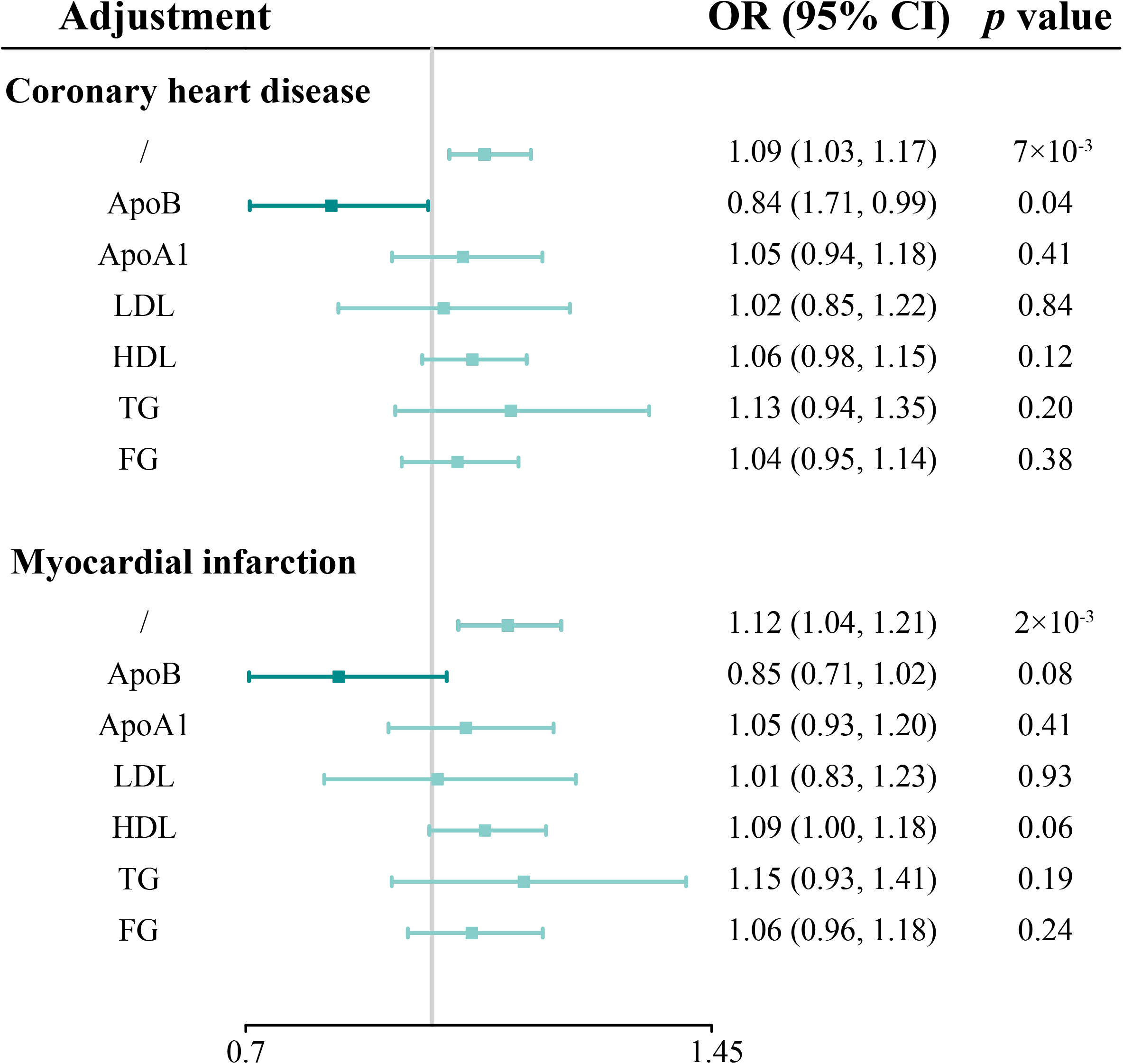
Multivariable Mendelian randomization analyses of genetically predicted IL-1Ra level with coronary heart disease and myocardial infarction on adjusting for cardiometabolic risk factors. Odds ratios are scaled per 1 standard deviation increase in the genetically predicted serum IL-1Ra level. IL-1Ra, interleukin 1 receptor antagonist; OR, odds ratio; CI, confidence interval.

**Figure 3.**
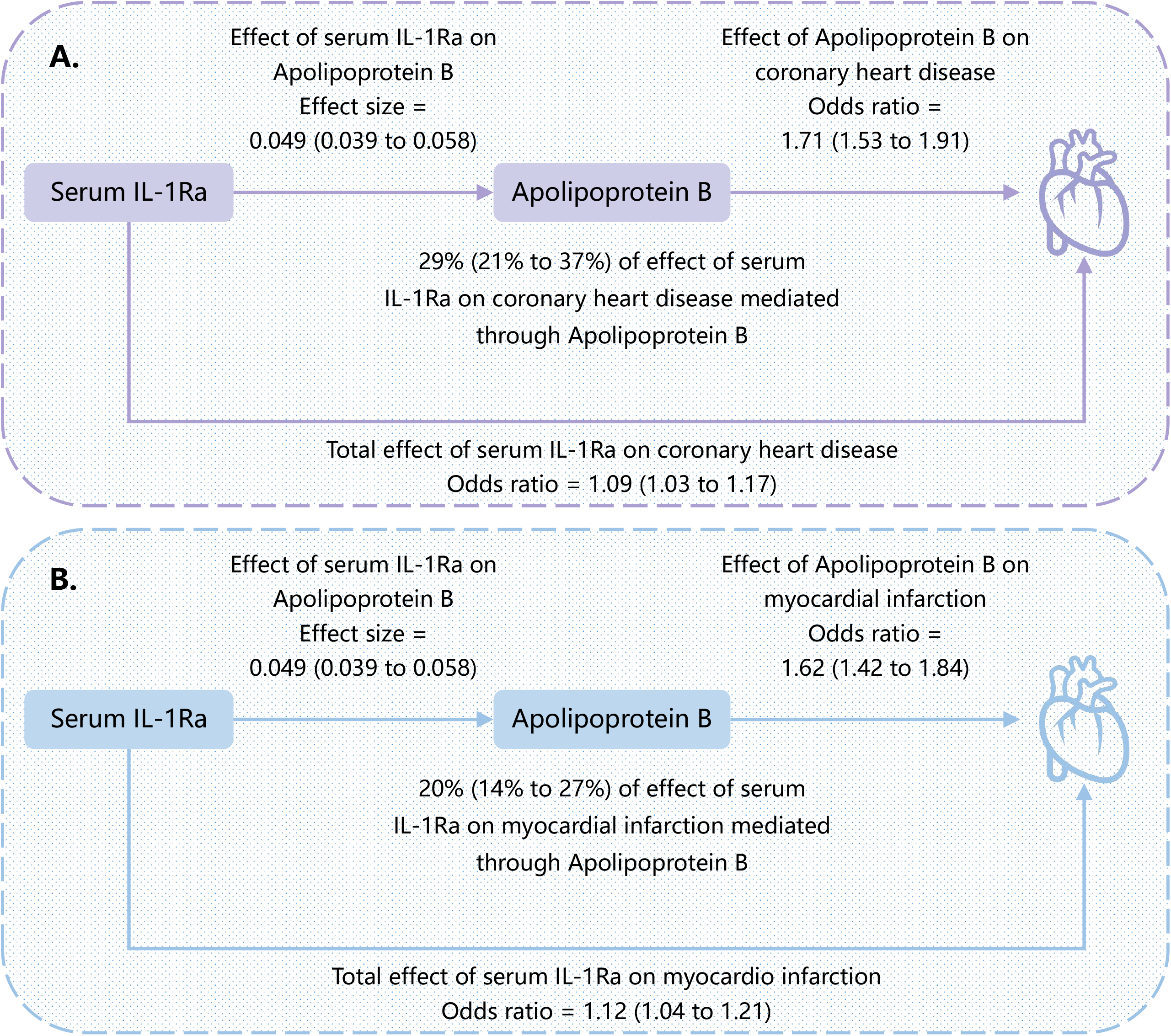
Causal directed acyclic graph showing the total effect of serum IL-1Ra on coronary heart disease and myocardial infarction, and the effect mediated by apolipoprotein B. The presented causal effect estimates with the corresponding 95% CI (shown in parenthesis) are scaled per one standard deviation increase in serum IL-1Ra. IL-1Ra, interleukin 1 receptor antagonist; CI, confidence interval.

We further investigated the associations of obesity and 11 lifestyle factors on serum IL-1Ra concentrations (Figure 4). Body mass index, waist circumference, VPA, sedentary behaviour, lifetime smoking index, and insomnia were positively associated with increased serum concentrations of IL-1Ra. On the other hand, sleep duration appeared to be inversely associated with IL-1Ra concentrations. No associations were detected for other studied lifestyle factors with IL-1Ra concentrations. In the mediation analysis, we found that adjusting for IL-1Ra concentrations led to a 47.8% attenuation of the causal effect estimate of sleep duration on the risk of CHD, and a 15.6% and 13.1% attenuation for smoking index and sedentary behaviour, respectively (Figure 5).

**Figure 4.**
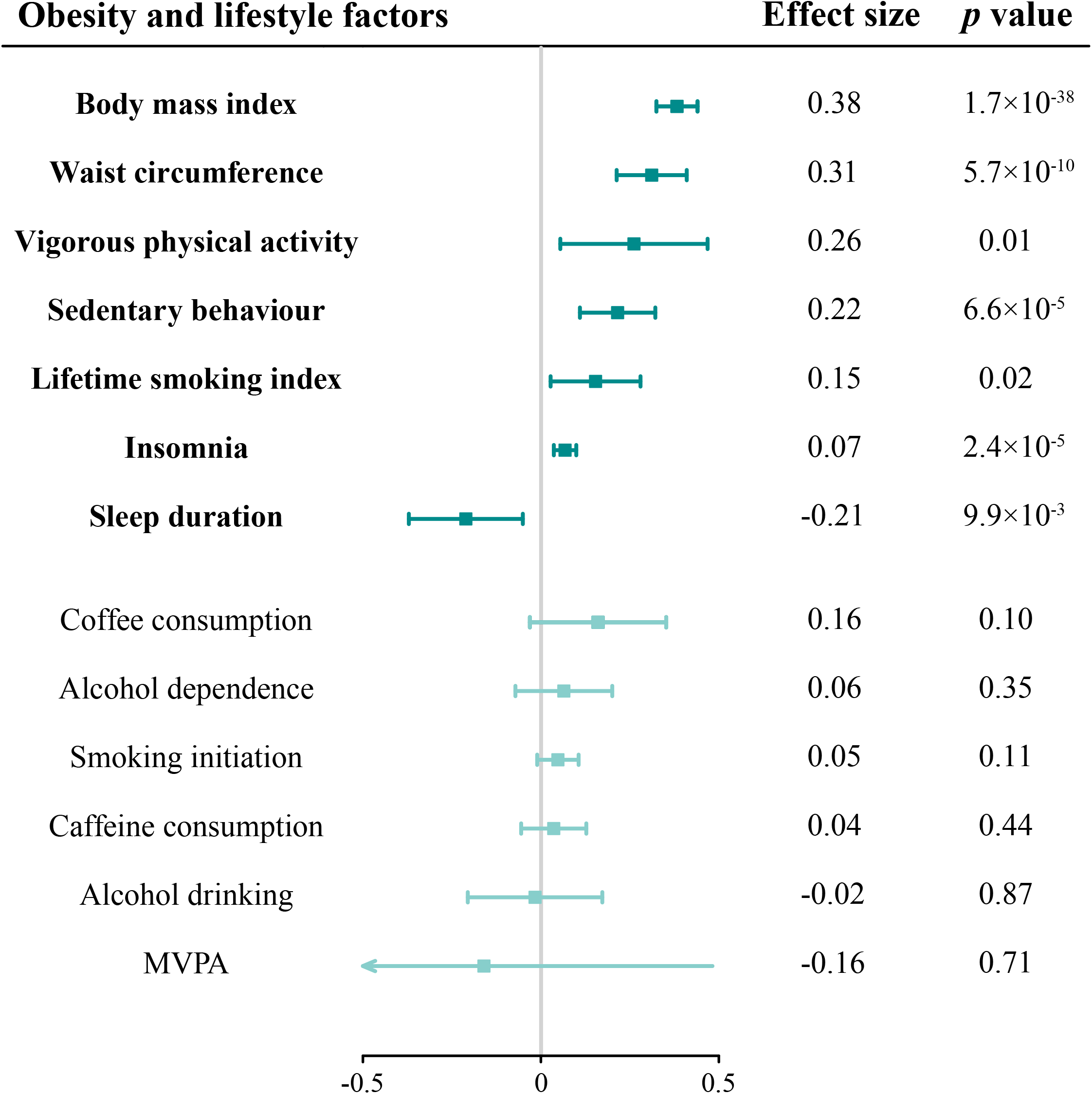
Causal associations of genetically predicted obesity and lifestyle factors with serum IL-1Ra level. IL-1Ra, interleukin 1 receptor antagonist.

**Figure 5.**
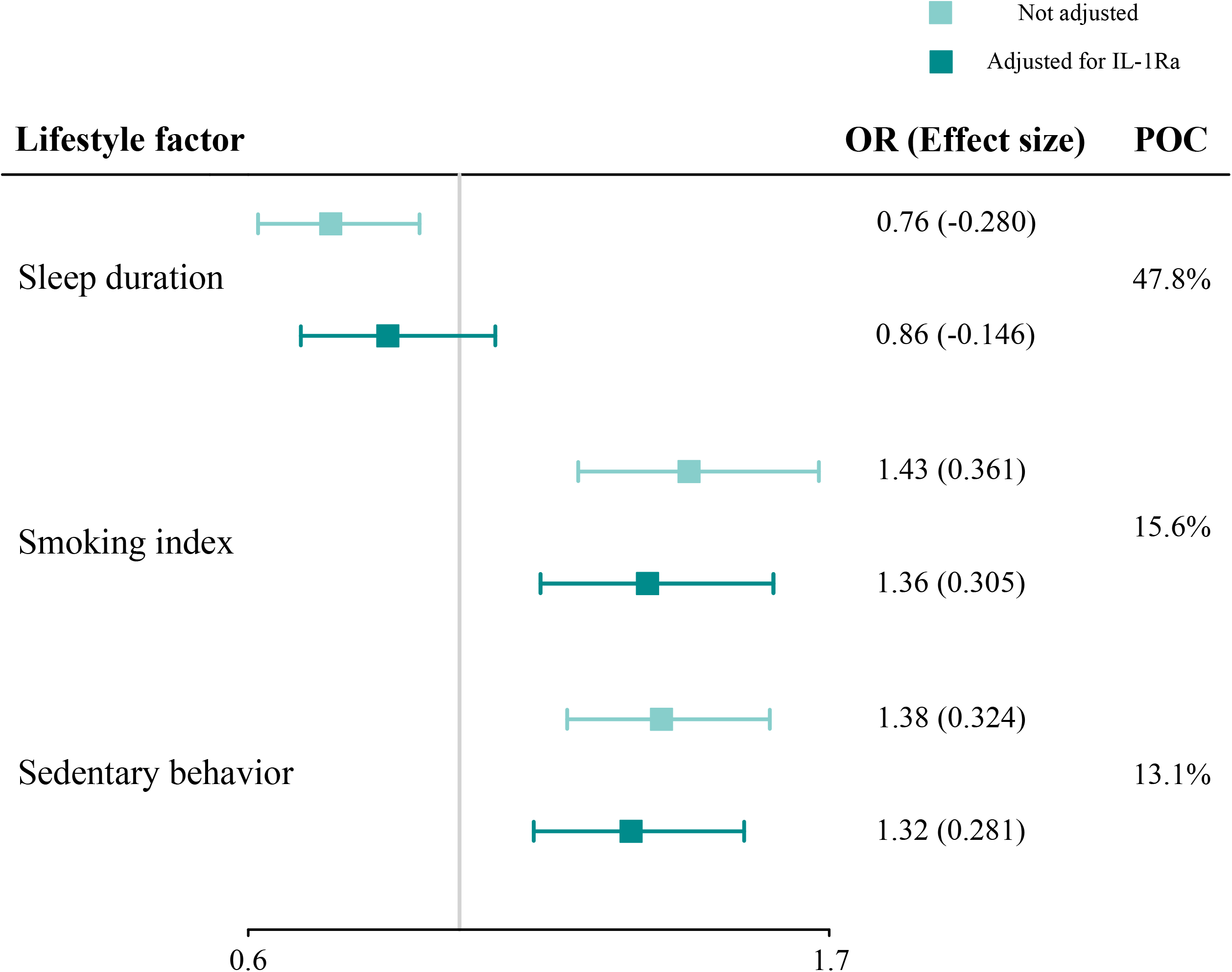
The causal effect of sleep duration, smoking index and sedentary behaviour on risk of coronary heart disease with or without adjustment for serum IL-1Ra level. IL-1Ra, interleukin 1 receptor antagonist; OR, odds ratio; POC, change of the proportion after adjusting for serum IL-1Ra level.

## Discussion

The current study employed a comprehensive framework of MR analyses based on large-scale human genetic data to investigate the causal associations of long-term IL-1 inhibition with the cardiometabolic profile. We found a significant positive association between serum IL-1Ra concentration and the risk of developing CHD and MI, as well as circulating levels of lipoprotein lipids, apolipoproteins and FG. Notably, the observed association between IL-1Ra and CHD/MI was primarily driven by apolipoprotein B. Several unhealthy lifestyle factors, like smoking, sedentary behaviour and insomnia were associated with increased concentrations of IL-1Ra. Mediation analysis suggested that IL-1Ra might mediate half of effects of a short sleep duration on CHD risk. These findings on the other side imply that in addition to drug treatment, appropriate lifestyle intervention helps to lower IL-1Ra concentration and consequently mitigate the risk of CHD.

Our findings were in line with a meta-analysis of six population-based cohort studies, involving over 20,000 participants, which revealed that the risk of CVD increased by about 11% (95% CI, 1.06-1.17), for every 1 SD increase in serum IL-1RA.^5^ Furthermore, the current study revealed the long-term and stable effect of lifetime exposure to elevated serum IL-1RA level. This observation was in agreement with our previous research, which demonstrated a causal association between IL-1RA concentration and CHD risk.^12^ Additionally, an early MR study suggested that long-term dual inhibition of IL-1α/β elevated LDL-cholesterol concentration, but not apolipoprotein or glycaemic traits.^13^ Nevertheless, our study provided consistent evidence for the causal effects of serum IL-1RA levels on apolipoprotein and FG. More importantly, we discovered that apolipoprotein B played a key role in mediating and potentially reversing the relationship between the IL-1RA and the risk of developing CHD, which may pave the way for new approaches to preventing and treating this condition.

The role of IL-1RA in incidence and development of CHD is not fully understood and remains controversial. On one hand, IL-1RA is known to bind to the IL-1 receptor, thereby preventing the binding of both IL-1α and IL-1β, and it has been proposed that inhibiting the IL-1 pathway may prevent the development of various CVDs, including atherosclerosis, MI, and heart failure.^7,41^ On the other hand, it has been indicated that IL-1 can directly affect lipid metabolism by suppressing the activity of lipoprotein lipase.^6^ Observational studies revealed a positive and significant correlation between IL-1RA and several molecules related to lipid metabolism, including cholesterol, triglycerides, and apolipoproteins B, all of which are known risk factors for CHD.^5,6^ Apolipoprotein B has been postulated as a key factor in the development of atherosclerosis, likely through the “response to retention” hypothesis.^42^ Lipoprotein particles, particularly those containing apolipoprotein B, become trapped in the innermost layer of arterial walls. The size and composition of apolipoprotein B particles, as well as the number of particles present in the bloodstream, would influence the tendency to be trapped in the arterial wall and contribute to the development of atherosclerosis.^42^ Genetic evidence also suggested that apolipoprotein B was the predominant lipids trait and may play a dominant role in the development of CHD.^19^ The effect estimates of other blood lipids were substantially attenuated, when adjusting for apolipoprotein B.^19^

Besides, the IL-1 receptor is widely distributed across human cells and plays crucial roles in numerous physiological processes, including host defence, wound healing, and autoimmunity, to name a few.^4^ Both IL-1α and IL-1β have been shown to have potentially beneficial cardiovascular effects, which might be hindered by the presence of IL-1Ra. Interestingly, an investigation conducted on an IL-1 receptor-deficient mouse indicated the cardioprotective effects of IL-1 signalling in advanced atherosclerosis.^43^ IL-1α, expressed by various cell types, not only mediates inflammation-related functions but also serves as an autocrine growth factor. IL-1β has also been shown to promote atheroprotective changes in advanced atherosclerotic lesions, such as facilitating outward remodelling and aiding in the formation and maintenance of the fibrous cap rich in smooth muscle cells and collagen.^44^

The link between lifestyle and the IL-1 pathway in CVDs is compelling. Emerging evidence suggests that an unhealthy lifestyle, such as a diet high in fat and calories, physical inactivity, sleep deficit and chronic stress, can markedly upregulate IL-1 production, triggering a cascade of pro-inflammatory responses.^45–47^ Sleep deprivation has been found to upregulate the expression and production of IL-1β in circulation and peripheral tissues across a variety of species, including humans, mice, rats, rabbits, cats, and monkeys, which underscores the critical role of sleep in regulating immune responses and inflammatory signaling pathways.^46^ Growing evidence suggests that regular physical activity exerts an overall anti-inflammatory effect by stimulating the release of muscle-derived myokines that enhance the production of IL-1Ra and IL-10, reducing dysfunction in adipose tissue and enhancing oxygen delivery.^47^ Cigarette smoking has also been linked to a diverse array of changes in immune and inflammatory markers, including IL-1RA, IL-1β and CRP, particularly among older individuals with a history of prolonged smoking.^48^

The present study employed a comprehensive MR approach to investigate the causal relationship and underlying mechanism between IL-Ra and CHD. The study design enabled unbiased estimation of causal effects from observational data and sensitivity analyses. Moreover, the potential interaction between various lifestyle factor and serum IL-Ra was further investigated. However, certain limitations should be acknowledged. First, the possibility of horizontal pleiotropy cannot be entirely ruled out, although this was not detected in MR-Egger intercept test and MR-PRESSO analysis. Second, the current study was restricted to individuals of European ancestry. Thus, the generalizability of the results to other populations may be limited. Third, the magnitude and duration of inhibition may differ between genetic and pharmacological IL-1 inhibition, and the findings need to be further investigated. Furthermore, the current study was limited to the use of summary-level data, and therefore, an investigation into the potential differences in the association patterns between sexes was not possible due to the lack of available sex-specific data.

## Conclusion

In summary, this study yields compelling evidence for the causal link between genetically predicted serum IL-1Ra and elevated risk of CHD/MI, with apolipoprotein B as the key driver, and a potential target for reversal. The potential cardiovascular benefits of a combined therapy involving IL-1 inhibition and lipid-modifying treatment aimed at apolipoprotein B merit further exploration, especially in individuals received IL-1Ra treatment, such as patients with RA. IL-1Ra appeared to mediate the associations between unhealthy lifestyle factors and CHD risk, which implies that lowering the levels of IL-1Ra among individuals with an unhealthy lifestyle may help lowering CHD risk.

## Acknowledgements

The authors thank all the investigators of corresponding genetic consortia and GWAS for providing the data publicly.

## Data Availability Statement

All the data used in the present study had been publicly available. The original contributions presented in the study are included in the article/Supplementary Material, further inquiries can be directed to the corresponding author/s.

## Author Contributions

SY, FY, NH and HC contributed to the conception or design of the work. FY, NH and SY contributed to the acquisition, analysis, or interpretation of data for the work. FY, NH and JS wrote the first draft of the manuscript with critical revisions from PS, LZ, SY and HC. All gave final approval and agree to be accountable for all aspects of work ensuring integrity and accuracy.

## References

1. Shaya GE, Leucker TM, Jones SR, et al. Coronary heart disease risk: Low-density lipoprotein and beyond. Trends Cardiovasc Med 2022; 32: 181–194.

2. Liberale L, Badimon L, Montecucco F, et al. Inflammation, Aging, and Cardiovascular Disease: JACC Review Topic of the Week. J Am Coll Cardiol 2022; 79: 837–847.

3. Abbate A, Toldo S, Marchetti C, et al. Interleukin-1 and the Inflammasome as Therapeutic Targets in Cardiovascular Disease. Circ Res 2020; 126: 1260–1280.

4. Mantovani A, Dinarello CA, Molgora M, et al. Interleukin-1 and Related Cytokines in the Regulation of Inflammation and Immunity. Immunity 2019; 50: 778–795.

5. Herder C, de Las Heras Gala T, Carstensen-Kirberg M, et al. Circulating Levels of Interleukin 1-Receptor Antagonist and Risk of Cardiovascular Disease: Meta-Analysis of Six Population-Based Cohorts. Arterioscler Thromb Vasc Biol 2017; 37: 1222–1227.

6. Almeida-Santiago C, Quevedo-Abeledo JC, Hernández-Hernández V, et al. Interleukin 1 receptor antagonist relation to cardiovascular disease risk in patients with rheumatoid arthritis. Sci Rep 2022; 12: 13698.

7. Dimosiari A, Patoulias D, Kitas GD, et al. Do Interleukin-1 and Interleukin-6 Antagonists Hold Any Place in the Treatment of Atherosclerotic Cardiovascular Disease and Related Co-Morbidities? An Overview of Available Clinical Evidence. J Clin Med 2023; 12: 1302.

8. Ikonomidis I, Tzortzis S, Andreadou I, et al. Increased benefit of interleukin-1 inhibition on vascular function, myocardial deformation, and twisting in patients with coronary artery disease and coexisting rheumatoid arthritis. Circ Cardiovasc Imaging 2014; 7: 619–628.

9. Morton AC, Rothman AMK, Greenwood JP, et al. The effect of interleukin-1 receptor antagonist therapy on markers of inflammation in non-ST elevation acute coronary syndromes: the MRC-ILA Heart Study. Eur Heart J 2015; 36: 377–384.

10. Burgess S, Davey Smith G, Davies NM, et al. Guidelines for performing Mendelian randomization investigations. Wellcome Open Res 2019; 4: 186.

11. Sekula P, Del Greco M F, Pattaro C, et al. Mendelian Randomization as an Approach to Assess Causality Using Observational Data. J Am Soc Nephrol 2016; 27: 3253–3265.

12. Yuan S, Lin A, He Q, et al. Circulating interleukins in relation to coronary artery disease, atrial fibrillation and ischemic stroke and its subtypes: A two-sample Mendelian randomization study. International Journal of Cardiology 2020; 313: 99–104.

13. Cardiometabolic effects of genetic upregulation of the interleukin 1 receptor antagonist: a Mendelian randomisation analysis. Lancet Diabetes Endocrinol 2015; 3: 243–253.

14. Folkersen L, Gustafsson S, Wang Q, et al. Genomic and drug target evaluation of 90 cardiovascular proteins in 30,931 individuals. Nat Metab 2020; 2: 1135–1148.

15. Clarke L, Zheng-Bradley X, Smith R, et al. The 1000 Genomes Project: data management and community access. Nat Methods 2012; 9: 459–462.

16. Nikpay M, Goel A, Won H-H, et al. A comprehensive 1000 Genomes–based genome-wide association meta-analysis of coronary artery disease. Nature Genetics 2015; 47: 1121–1130.

17. Okada Y, Wu D, Trynka G, et al. Genetics of rheumatoid arthritis contributes to biology and drug discovery. Nature 2014; 506: 376–381.

18. Ligthart S, Vaez A, Võsa U, et al. Genome Analyses of >200,000 Individuals Identify 58 Loci for Chronic Inflammation and Highlight Pathways that Link Inflammation and Complex Disorders. Am J Hum Genet 2018; 103: 691–706.

19. Richardson TG, Sanderson E, Palmer TM, et al. Evaluating the relationship between circulating lipoprotein lipids and apolipoproteins with risk of coronary heart disease: A multivariable Mendelian randomisation analysis. PLOS Medicine 2020; 17: e1003062.

20. Chen J, Spracklen CN, Marenne G, et al. The trans-ancestral genomic architecture of glycemic traits. Nat Genet 2021; 53: 840–860.

21. Evangelou E, Warren HR, Mosen-Ansorena D, et al. Genetic analysis of over 1 million people identifies 535 new loci associated with blood pressure traits. Nat Genet 2018; 50: 1412–1425.

22. Pulit SL, Stoneman C, Morris AP, et al. Meta-analysis of genome-wide association studies for body fat distribution in 694 649 individuals of European ancestry. Hum Mol Genet 2019; 28: 166–174.

23. Shungin D, Winkler TW, Croteau-Chonka DC, et al. New genetic loci link adipose and insulin biology to body fat distribution. Nature 2015; 518: 187–196.

24. Liu M, Jiang Y, Wedow R, et al. Association studies of up to 1.2 million individuals yield new insights into the genetic etiology of tobacco and alcohol use. Nat Genet 2019; 51: 237–244.

25. Wootton RE, Richmond RC, Stuijfzand BG, et al. Evidence for causal effects of lifetime smoking on risk for depression and schizophrenia: a Mendelian randomisation study. Psychol Med 2020; 50: 2435–2443.

26. Walters RK, Polimanti R, Johnson EC, et al. Transancestral GWAS of alcohol dependence reveals common genetic underpinnings with psychiatric disorders. Nat Neurosci 2018; 21: 1656–1669.

27. Zhong VW, Kuang A, Danning RD, et al. A genome-wide association study of bitter and sweet beverage consumption. Hum Mol Genet 2019; 28: 2449–2457.

28. Cornelis MC, Kacprowski T, Menni C, et al. Genome-wide association study of caffeine metabolites provides new insights to caffeine metabolism and dietary caffeine-consumption behavior. Hum Mol Genet 2016; 25: 5472–5482.

29. Klimentidis YC, Raichlen DA, Bea J, et al. Genome-wide association study of habitual physical activity in over 377,000 UK Biobank participants identifies multiple variants including CADM2 and APOE. Int J Obes (Lond) 2018; 42: 1161–1176.

30. van de Vegte YJ, Said MA, Rienstra M, et al. Genome-wide association studies and Mendelian randomization analyses for leisure sedentary behaviours. Nat Commun 2020; 11: 1770.

31. Dashti HS, Jones SE, Wood AR, et al. Genome-wide association study identifies genetic loci for self-reported habitual sleep duration supported by accelerometer-derived estimates. Nat Commun 2019; 10: 1100.

32. Jansen PR, Watanabe K, Stringer S, et al. Genome-wide analysis of insomnia in 1,331,010 individuals identifies new risk loci and functional pathways. Nat Genet 2019; 51: 394–403.

33. Burgess S, Butterworth A, Thompson SG. Mendelian Randomization Analysis With Multiple Genetic Variants Using Summarized Data. Genetic Epidemiology 2013; 37: 658–665.

34. Bowden J, Davey Smith G, Haycock PC, et al. Consistent Estimation in Mendelian Randomization with Some Invalid Instruments Using a Weighted Median Estimator. Genet Epidemiol 2016; 40: 304–314.

35. Bowden J, Davey Smith G, Burgess S. Mendelian randomization with invalid instruments: effect estimation and bias detection through Egger regression. Int J Epidemiol 2015; 44: 512–525.

36. Verbanck M, Chen C-Y, Neale B, et al. Detection of widespread horizontal pleiotropy in causal relationships inferred from Mendelian randomization between complex traits and diseases. Nat Genet 2018; 50: 693–698.

37. Burgess S, Thompson SG. Multivariable Mendelian randomization: the use of pleiotropic genetic variants to estimate causal effects. Am J Epidemiol 2015; 181: 251–260.

38. Burgess S, Daniel RM, Butterworth AS, et al. Network Mendelian randomization: using genetic variants as instrumental variables to investigate mediation in causal pathways. Int J Epidemiol 2015; 44: 484–495.

39. Hemani G, Zheng J, Elsworth B, et al. The MR-Base platform supports systematic causal inference across the human phenome. eLife 2018; 7: e34408.

40. Yavorska OO, Burgess S. MendelianRandomization: an R package for performing Mendelian randomization analyses using summarized data. Int J Epidemiol 2017; 46: 1734–1739.

41. Herder C, Donath MY. Interleukin-1 receptor antagonist: friend or foe to the heart? The Lancet Diabetes &Endocrinology 2015; 3: 228–229.

42. Sniderman AD, Thanassoulis G, Glavinovic T, et al. Apolipoprotein B Particles and Cardiovascular Disease: A Narrative Review. JAMA Cardiol 2019; 4: 1287–1295.

43. Alexander MR, Moehle CW, Johnson JL, et al. Genetic inactivation of IL-1 signaling enhances atherosclerotic plaque instability and reduces outward vessel remodeling in advanced atherosclerosis in mice. J Clin Invest 2012; 122: 70–79.

44. Gomez D, Baylis RA, Durgin BG, et al. Interleukin-1β has atheroprotective effects in advanced atherosclerotic lesions of mice. Nat Med 2018; 24: 1418–1429.

45. Pavillard LE, Marín-Aguilar F, Bullon P, et al. Cardiovascular diseases, NLRP3 inflammasome, and western dietary patterns. Pharmacol Res 2018; 131: 44–50.

46. Zielinski MR, Gibbons AJ. Neuroinflammation, Sleep, and Circadian Rhythms. Frontiers in Cellular and Infection Microbiology; 12, https://www.frontiersin.org/articles/10.3389/fcimb.2022.853096 (2022, accessed 17 April 2023).

47. Nieman DC, Wentz LM. The compelling link between physical activity and the body’s defense system. J Sport Health Sci 2019; 8: 201–217.

48. Shiels MS, Katki HA, Freedman ND, et al. Cigarette smoking and variations in systemic immune and inflammation markers. J Natl Cancer Inst 2014; 106: dju294.

